# Leveraging genome-enabled growth models to study shoot growth responses to water deficit in rice

**DOI:** 10.1101/690479

**Authors:** Malachy Campbell, Alexandre Grondin, Harkamal Walia, Gota Morota

## Abstract

Elucidating genotype-by-environment interactions (G×E) and partitioning its contribution to the phenotypic variation remains a long standing challenge for plant scientists. Recent quantitative genetic frameworks have improved predictions of G× E responses. However, these models do not explicitly consider the processes that give rise to G×E. To overcome this limitation, we propose a novel framework to elucidate the genetic basis of dynamic shoot growth trajectories under contrasting water regimes using genome-wide markers to model genotype-specific shoot growth trajectories as a function of soil water availability. A rice diversity panel was phenotyped daily over a period of twenty-one days during the early vegetative stage using an automated, high-throughput image-based, phenotyping platform that enabled us to estimate daily shoot biomass and soil water content. Using these data, we modeled shoot growth as a function of time and soil water content, and were able to determine the soil water content and/or time point where an inflection in the growth trajectory occurred. We found that larger, more vigorous plants tend to exhibit an earlier repression in growth compared to smaller, slow growing plants, indicating a potential trade off between early vigor and tolerance to prolonged water deficits. We integrated the growth model within a hierarchical Bayesian framework and used marker information to estimate model parameters and the associated loci through genome-wide association analysis. Genomic inference for model parameters and time of inflection (TOI) identified several candidate genes. Among them an aquaporin, *OsPIP1;1* was identified as a candidate for time of inflection under drought and showed significantly lower expression in accessions exhibiting later TOI in drought. This study is the first to utilize a genome-enabled growth model to study drought responses in rice, and presents a new approach to jointly model dynamic morpho-physiological responses and environmental covariates.

## Introduction

Rice is one of the most important food crops, and is a major source of food security for more than 3.5 billion people worldwide. Adequate water availability is essential for proper vegetative growth and grain development. Approximately 40 million hectares of rainfed rice is grown worldwide, with the majority of production being concentrated in developing nations (Singh and Singh, 2000). Erratic precipitation events, as well as the increased competition for fresh water for non-agricultural uses has become a major constraint for rice production (Korres et al., 2017).

Given the socioeconomic impacts of water limitations, improving drought tolerance is a major target for breeding programs. However, the multiple and often unpredictable drought stress scenarios in drought prone environment makes improvement of drought tolerance in rice challenging. Further, traits that are important for adaptation to limited water availability, particularly morpho-physiological traits, are complex and often have low heritability (Kamoshita et al., 2008). These characteristics impede the discovery of loci that confer large effects on the phenotype, and limit the utility of marker-assisted selection for improving drought tolerance.

Recent advances in phenomics and genomics have offered new tools for discovering and quantifying traits associated with drought adaptation, and their genetic basis (Berger et al., 2010; Furbank and Tester, 2011; Araus and Cairns, 2014). Access to high-throughput, image-based phenomic systems in the public sector has allowed researchers to non-destructively measure traits of interest for large populations in highly controlled greenhouse or field environments. These data can be leveraged to uncover complex physiological responses to suboptimal conditions throughout the growing season and understand their contribution to agronomic performance. Several studies have used these traits as covariates in the conventional genomic prediction frameworks to increase prediction accuracies for agronomic traits such as yield (Aguate et al., 2017; Montesinos-López et al., 2017; Sun et al., 2017; Krause et al., 2019). However, with these approaches it is unclear how these secondary phenotypes are contributing to trait of interest.

Process-based eco-physiological models seek to predict outcomes by explicitly modeling the interaction of biological processes with environmental covariates (Batchelor et al., 2002; van Ittersum et al., 2003; Parent and Tardieu, 2014). These models are routinely used to predict the development or productivity of a crop in a given set of environments. However, a major disadvantage of these models is that genotypic variation is often unaccounted or not optimally utilized in the predictions (Onogi et al., 2016). Thus, their application in genomic prediction or inference studies is limited. Several studies have sought to integrate crop growth models with established quantitative genetic frameworks (Technow et al., 2015; Onogi et al., 2016; Wang et al., 2019). For instance, Technow et al. (2015) used an approximate Bayesian computation framework to integrate crop growth modeling and whole-genome prediction to predict yield in maize. The authors showed a clear advantage of the genome-enabled crop growth model over the conventional genomic prediction approach using simulated data. More recently, Onogi et al. (2016) leveraged a crop growth model to predict heading date in rice. The authors integrated the phenological model proposed by Yin et al. (1997) and implemented by Nakagawa et al. (2005) with a whole-genome prediction using a hierarchical Bayesian approach. The hierarchical Bayesian approach outperformed conventional genomic prediction models as well as approaches that fit the crop growth model and genomic prediction model in separate steps. The advantage of the integrated approaches proposed by Technow et al. (2015) and Onogi et al. (2016) is that model parameter estimates are informed by the genomic relationships among the accessions, which can improve the accuracy of the parameter estimates. Moreover, since these approaches are based on a Bayesian whole-genome regression framework, it predicts markers effects, enabling marker level association with model parameters. However, to date no studies have leveraged these genome-enabled crop growth models for biological inference or to elucidate the genetic loci that influence model parameters.

In the current study, we sought to leverage the frameworks developed by Onogi et al. (2016) to study the effects of water deficit on shoot growth trajectories for a diverse set of rice accessions. To this end, rice accessions were subjected to drought stress (20 % field capacity, FC) and shoot growth was quantified over 20 days using an image-based phenomics platform. A corresponding set of rice accessions was maintained under optimal water conditions (90 % FC). The automated phenotyping system allowed us to estimate daily water use by each accession and the soil water content. Together, these data were used to develop a novel growth model that models shoot growth trajectories as a function of soil water content and time. This growth model was integrated into the hierarchical Bayesian framework of Onogi et al. (2016) to elucidate the genes underlying model parameters. This approach provides a biologically meaningful framework that simultaneously (1) models the interrelationship between growth rate and soil water availability, (2) estimates quantitative trait loci (QTL) effects for model parameters, and (3) provides genetic values for model parameters that can be used for genetic evaluation.

## Materials and Methods

### Plant materials and greenhouse conditions

A subset of the Rice Diversity Panel 1 were used in this study (Zhao et al., 2011). Seed preparation was performed following Campbell et al. (2015). Briefly, seeds were surface sterilized with Thiram fungicide and were germinated on moist paper towels in plastic boxes for 3 days. Three uniform seedlings were selected and transplanted to pots (150mm diameter x 200 mm height) filled with approximately 2.5 kg of UC Mix. Square containers were placed below each pot to allow water to collect. Temperatures in the greenhouses were maintained at 28/26.0°C (day/night), and relative humidity was maintained at approximately 60% throughout the day and night.

### Experimental Design

Three hundred seventy-eight accessions were phenotyped at the Plant Accelerator, Australian Plant Phenomics Facility, at the University of Adelaide, SA, Australia in three independent experiments. The experiments were performed from February to April 2016. The 378 accessions were randomly partitioned in two smarthouses, each of which consisted of 432 pots positioned across 24 lanes. In each experiment, a subset of 54 accessions were randomly selected and two replicates were planted for each accession/treatment combination. A split-plot design was employed with two consecutive pots having the same accession, but with the two different water regimes randomly assigned to them.

Seven days after transplanting to soil, plants were thinned to one seedling per pot and two layers of blue mesh was placed on top of the soil to reduce soil water evaporation. At 11 days after transplant (DAT), the plants were loaded on the imaging system and were watered to 90% field capacity. Water was withheld from one of the two pots for each accession beginning at 13 DAT. Water was withheld until the end of the experiment or until the FC reached 20%, after which the plants were maintained at 20% FC.

### Image analysis

The plants were imaged each day from 13 to 33 DAT using a visible (red–green–blue camera; Basler Pilot piA2400–12 gc, Ahrensburg, Germany) from two side-view angles separated by 90 degree and a single top view. The LemnaGrid software was used to extract “plant pixels” from RGB images. The image analysis pipeline is identical to that described in Campbell et al. (2018). “Plant pixels” from each of the RGB images for each plant and time point were summed and was used as a proxy for shoot biomass. We refer to this digital trait as projected shoot area (PSA). Several studies have shown that this metric is accurate representation of shoot biomass (Campbell et al., 2015; Golzarian et al., 2011; Knecht et al., 2016). Outlier plants at each time point were detected for each trait using the 1.5(IQR) rule. Plants that were flagged as potential outliers were plotted and inspected visually and those that exhibited abnormal growth patterns were removed prior to downstream analyses. In total, 221 plants were removed, leaving 2,586 plants for downstream analyses. Since the genome-enabled crop growth model does not accommodate missing data, accessions with missing values were excluded from further analyses. This culling resulted in a total of 349 accessions being used for downstream analyses.

### Modeling shoot growth as a function of time and soil water content

To model the effects of water deficit on shoot growth trajectories, we devised a growth model that is essentially an extension of the classical Gompertz growth model. The Gompertz growth model was modified so that shoot growth trajectories were modeled as a function of time and soil water content. This model is referred to as the WSI-Gomp model in the remainder of the manuscript. The WSI-Gomp model and its relationship with the classical Gompertz growth model are discussed in greater detail in the Results section (see “Defining the growth model”). The WSI-Gomp model is given by

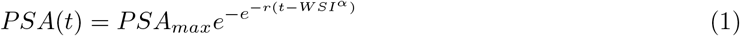

where *PSA_max_* is a parameter that describes the maximum biomass achieved by the plant; *r* is a parameter that describes the absolute growth rate; *t* is a vector of standardized time values [0,1]; and *α* is a genotype-specific tuning parameter that modifies the effect of WSI on PSA. WSI is the water stress index, a unitless index that describes the severity of water stress, and is given by

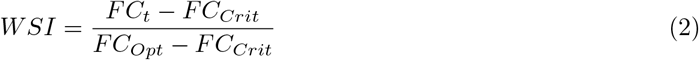

*FC_t_* is the portion of field capacity at time *t*; *FC_Crit_* (critical field capacity) is the proportion of field capacity in which growth ceases; *FC_Opt_* is the proportion of field capacity that is optimal for growth. *FC* was calculated at each time point from pot weights given by the automated watering system. Since *FC_Crit_* and *FC_Opt_* are unknown and likely to be genotype-dependant, we assumed that the optimal conditions for growth in rice occur when the soil is completely saturated, and the critical value for FC is equal to 0.1. Although, these assumptions require empirical evidence to validate, they provide a standardized metric that describes soil water content in a decreasing non-linear trend that is on the same scale as the standardized time values. These characteristics allow PSA to be modeled as a function of time and soil water content using the Gompertz growth model.

### Leveraging whole genome regression to estimate model parameters

The “integrated approach” developed by Onogi et al. (2016) uses a hierarchical Bayesian framework to simultaneously infer of growth model parameters and marker effects. Thus, by leveraging the genetic relationships between accessions, the integrated approach should yield more accurate solutions for the model parameters. The details of the “integrated approach” is given in Onogi et al. (2016). Briefly, solutions for the growth model parameters are regressed on genome-wide markers and extended Bayesian LASSO (EBL) is used to predict marker effects for each of the model parameters. The regression model is given by

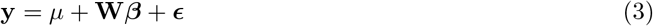

**W** is a *n* × *m* matrix of marker genotypes coded as −1, 0, 1 and *n* is the number of accessions (349) and *m* is the number of markers (33,697); *μ* is the intercept for each parameter; *β* is a *m* × 1 vector of predicted marker effect for each model parameter. The prior distribution of marker effects for marker *i* is

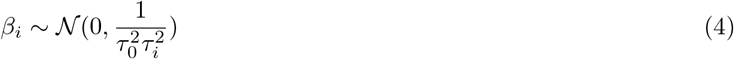

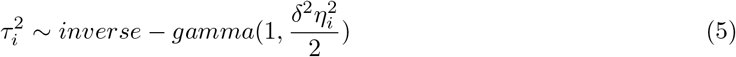

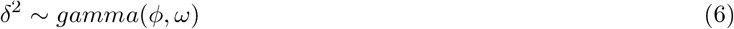

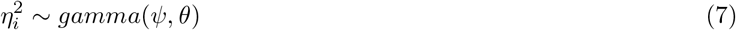

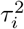 is the precision for the effect of marker *i*; 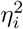 is the marker-specific shrinkage parameter for marker *i*; *δ*^2^ is the global shrinkage parameter; and *ω*, *ϕ*, *θ*, and *ψ* are hyperparameters. Default values were used for hyperparameters. We assume the following for WSI-Gomp model parameters

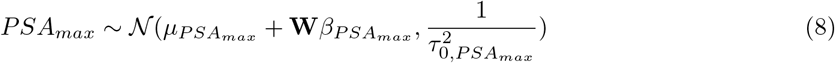

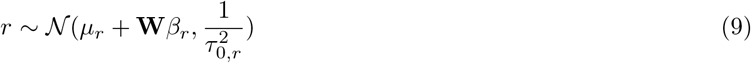

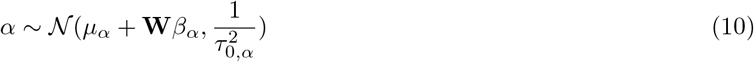

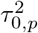 is the residual precisions for model parameter *p*. Moreover, for the residuals we assume 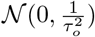. With the “integrated approach” proposed by Onogi et al. (2016), model parameters are inferred using a variational Bayes approach in which means and variances of the growth model parameters are obtained using Markov chain Monte Carlo sampling and are used to update EBL parameters.

### Genome-wide association for time of inflection

We sought to utilize the WSI-Gomp model to identify genomic loci that influenced the timing of the transition to a declining growth rate. To this end we used model parameters obtained from the hierarchical Bayesian approach described above and observed WSI values to solve for the time of inflection (TOI). In the classical Gompertz growth model, 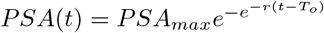, the growth rate begins to decline when the (*t* – *T_o_*) term becomes positive (e.g. when t exceeds *T_o_*). Thus, *T_o_* can be defined as the time of inflection. In the WSI-Gomp model, this component is given by (*t* – *WSI^α^*). Thus, the time of inflection occurs when t ≥ *WSI^α^*. Using the hierarchical Bayesian approach, we obtained estimates for α for each accession in drought and control conditions, and used these values to solve for TOI using WSI values for each corresponding plant in each experiment. TOI was defined as the first day in which (*t* – *WSI^α^*) was positive. This yielded a single TOI value for each plant in each experiment.

These TOI values were used as a derived phenotype for further genome-wide association study (GWAS) analysis. The following Bayesian LASSO regression model was fit using the BGLR package (Pérez and de Los Campos, 2014)

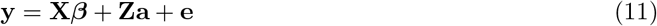

where **X** is an incidence matrix relating the vector ***β*** of fixed effects for experiment to observations, **Z** is an incidence matrix relating the vector of random marker effects **a** to **y**, and **e** is the residual. Since the vector **y** is a vector of discrete TOI values, **y** was treated as an ordinal response and a probit link function was used. In Bayesian LASSO, the marginal prior distribution for each marker effect is a double exponential function that includes an unknown parameter λ^2^ with a prior distribution λ^2^ ~ gamma(*r*, *s*) (Pérez and de Los Campos, 2014). BGLR sets *s* = 1.1 by default and solves for *r* based on the “prior” R^2^ of the model. Details on the BL approach implemented in BGLR is provided in the package vignette. A Gaussian prior with mean zero and variance equal to 1 × 10^10^ was used for fixed effects.

### Gene expression analysis

To assess the expression of candidate genes we utilized a publicly available data set consisting of RNA sequencing data for 91 diverse rice accessions (Campbell et al., 2019). Of these 91 accessions, 87 overlapped with accessions included in this study. The collection, processing and analysis of these data are described in detail in Campbell et al. (2019). Briefly, best linear unbiased prediction (BLUPs) were obtained for each gene and these values were transformed into the quantiles of a standard normal distribution with ties broken randomly. This ensured that the expression of each gene was normally distributed and inference from the linear model was reliable. To compare expression levels between allelic groups at a given SNP, a linear model was fit that included the first four principle components of the kinship matrix and the SNP genotype. All terms were considered fixed. This model was compared to a null model that included all terms described above, but omitted the SNP genotype. The two models were compared using a likelihood ratio test. The residuals from the null model were used to plot expression levels as shown in Figure 7. Thus these residuals show the expression level in each accession while accounting for population structure.

## Results

### Image-based phenotyping captures the sensitivity of rice to drought stress

To examine drought responses in rice (*Oryza sativa*), a diversity panel was phenotyped over a period of 21 days during the early vegetative stage using an automated high-throughput phenotyping platform (Supplemental File S1). The diversity panel consists of 349 accessions from 79 countries, and captures much of the genetic diversity within cultivated rice (Zhao et al., 2011). The 349 accessions were grown in a partially replicated paired design, in which for each accession the control, well watered plants were grown alongside the drought stressed plant.

All plants were watered to 90% FC nine days after transplanting (13 day-old plants), and water was withheld from day 14 on wards for the drought treatment plants. The drought stressed plants were only rewatered if their FC dropped below 20%. A simple *t*-test was carried out at each time point to determine when a significant reduction in soil water availability was experienced. A significant difference in pot water content (FC) was observed from second day of imaging (Figure 1; *p* < 0.0024, Bonferroni’s correction with *α* = 0.05), when the drought plants on average were at 90.9% FC. This time point was selected to mark the onset of drought stress.

**Figure 1.**
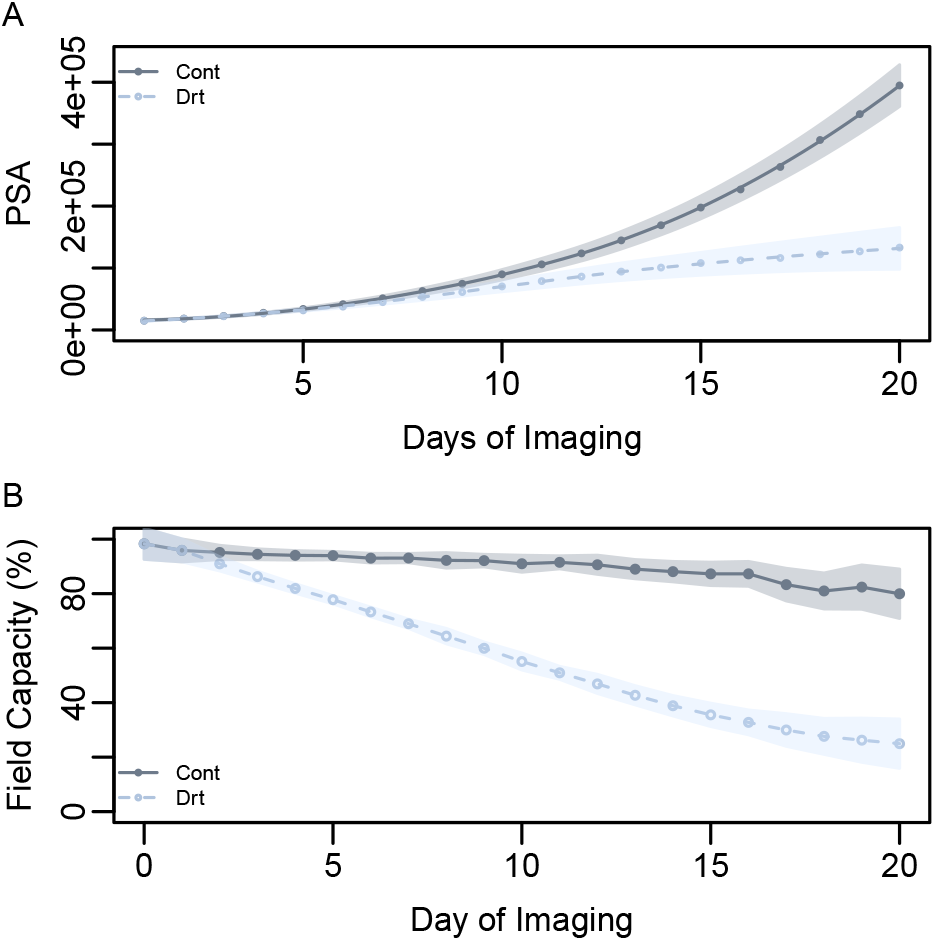
Effect of water deficit on shoot growth. (A) Mean shoot growth trajectories (PSA) in drought and control conditions over 21 days of imaging. (B) Mean percent field capacity in drought and control conditions over the 21 days of imaging. Water was withheld starting at day 1 of imaging. The shaded regions indicate the standard deviation for each treatment.

The impact of drought stress on shoot growth (biomass) was estimated from RGB images and expressed as a digital metric called PSA. An ANOVA was carried out at each time point using the following linear model *PSA_t_* = *μ_t_* + *Trt_a_* + *Acc_i_* + *Trt_a_* × *Acc_ai_* + *e_i_*, where *PSA_t_* is the trait at time *t* for accession *i* and treatment *a*, *μ* is the overall mean at time *t*, *Trt* is the effect of the *a^th^* treatment, *Acc_i_* is the effect of the *i^th^* accession, *Trt* × *Acc_ai_* is the interaction between *Trt* and *Acc*. Significant effects for the interaction between accession and treatment were observed from day 16 onward. Thus, during the early time points the main effects of treatment could be interpreted. Drought treatment had a significant effect on PSA beginning on the fourth day of imaging (Figure 1B; *p* < 0.0024, Bonferroni’s correction with *α* = 0.05). Interestingly, at this time point drought treated plants, on average, were at 81.9% FC, which is only approximately 12.5% below control plants. These data suggests that even a small limitation of water can have a significant impact on shoot growth in rice, and thus confirming the high level of drought sensitivity reported for rice (Lafitte et al., 2004).

### Defining the growth model

The Gompertz growth model has been used extensively to model asymptotic processes that exhibit a sigmoid trend (Winsor, 1932). For the drought conditions imposed in the current study, we expect shoot growth to follow an exponential trajectory during the initial time points when soil water is not limiting. However, as the soil dries out the growth rate should slow, and eventually when soil water content falls below some threshold, growth should cease completely. This sigmoid/asymptotic trend is to some degree visible in the mean growth trajectory in Figure 1A. Thus, the trend exhibited by plants in drought conditions can be modeled using the Gompertz growth model. The classical Gompertz model is given by 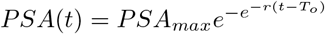, where *t* is a vector of time values, *r* is the absolute growth rate, *PSA_max_* is the maximum biomass (e.g. asymptote), and *T_o_* is the inflection point in the growth curve where the relative growth rate begins to slow. Figure 2A provides a graphical summary of the classical Gompertz model. While the classical Gompertz model provides an intuitive framework to model asymptotic growth trajectories, in its current form it does not accommodate for environmental data. Thus it cannot be used to address how shoot growth varies in response to soil water content.

To address this limitation, we sought to modify the Gompertz growth model so that shoot growth trajectories could be modeled as a function of time and soil water content. The objective was to develop a model with a similar form to the classical Gompertz model, but allowed us to determine the soil water content value that results in inflection of the growth trend. Since the automated phenotyping system provides daily records for soil water content for each plant, we defined an index (water stress index, WSI) that reflects the severity of water stress. WSI is given by 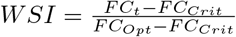, where *FC_t_* indicates the percent field capacity (FC) at time *t*, *FC_Opt_* is the optimal percent field capacity for growth, and *FC_Crit_* is the percent field capacity at which growth ceases. Since these values are expected to vary depending on the genotype, we assumed that growth will cease at 10% FC (*FC_Crit_* = 1) and the growth will proceed optimally when the soil is saturated (*FC_Opt_* = 100). This equation provides a unitless metric that will vary between 0 and 1, with higher values indicating lower water stress and lower values indicating a greater stress pressure. For this metric to be introduced into the Gompertz growth model, we standardized the time values so that they ranged from 0 to 1. Finally, we introduced a third parameter (*α* into the model that act as a genotype-dependant tuning parameter and modifies the effect of WSI on growth rate. This new WSI-integrated model (WSI-Gomp) is given by 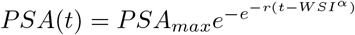. The WSI-Gomp model is shown in Figure 2B.

**Figure 2.**
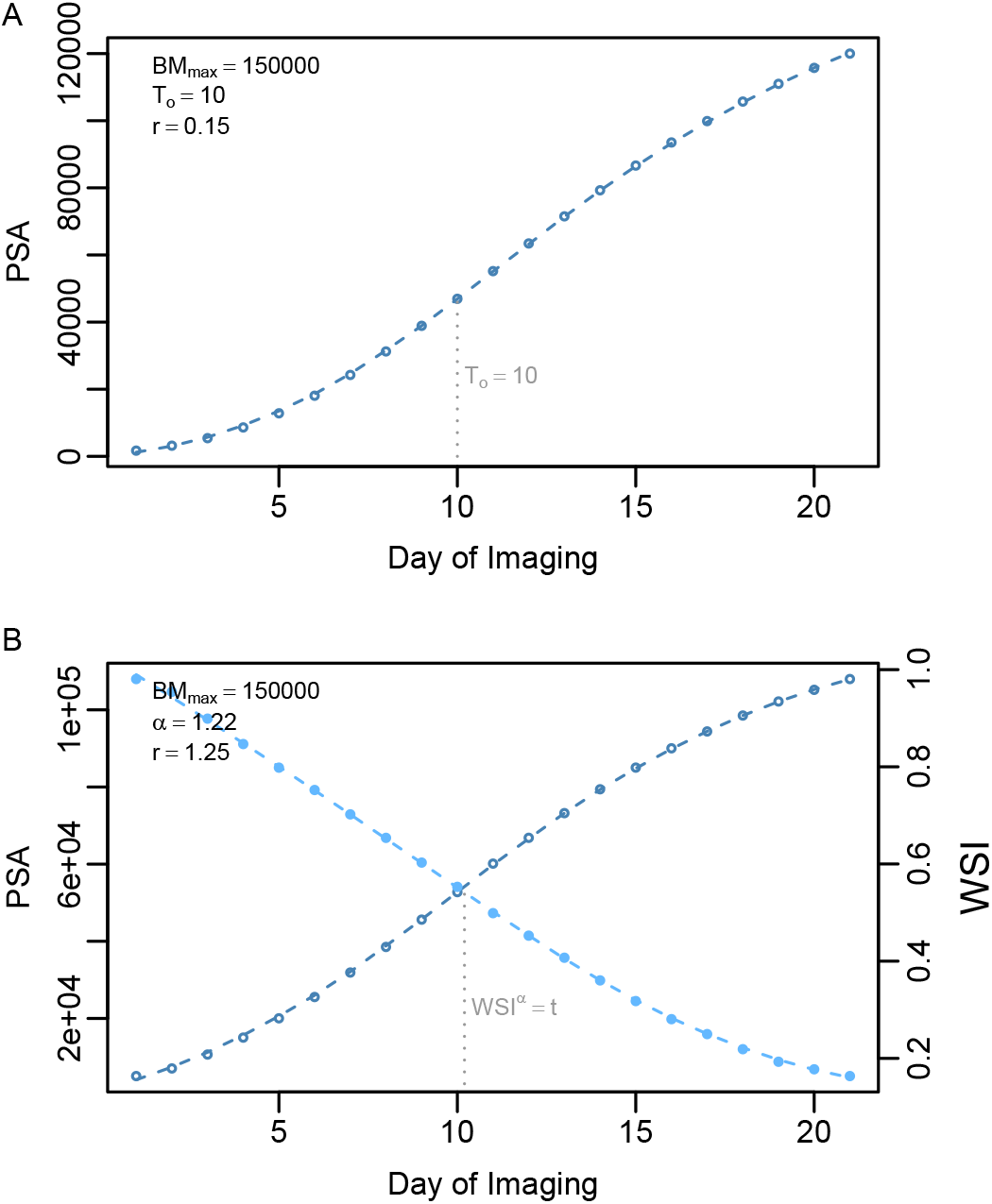
Graphical representation of the classical Gompertz model and the WSI-Gomp model. (A) The classical Gompertz growth model was used to generate PSA values over a 21-day period. The parameter values used are provided in the top left corner of the plot. The grey, vertical broken line indicates the inflection point (*T_o_*). (B) The WSI-Gompertz growth model was used to generate PSA values over a 21-day period. PSA values are shown using dark blue points and broken line. The light blue points and line indicate the WSI values over the 21-day period The grey, vertical broken line indicates the inflection point (*T_o_*).

To capture the effects of soil water deficit on growth trajectories, the WSI-Gomp model was fit to growth trajectories in drought and control conditions for each of the 349 accessions using a novel hierarchical Bayesian approach that leverages the genetic relationships among lines to obtain solutions for the model parameters (Onogi et al., 2016). Model parameter estimates for each accession were used to predict growth trajectories using observed WSI values. The ability of the WSI-Gomp model to capture these dynamic responses was assessed by comparing predicted PSA values and observed values at each time point using two metrics: root mean squared error (RMSE) and Pearson’s correlation. Overall, the WSI-Gomp model provided a good fit to the observed shoot growth trajectories (Figure 3). The correlation between observed and predicted PSA values ranged from 0.41 to 0.87 in control. While the correlation was slightly lower in drought conditions and ranged from 0.52 to 0.75. Correlation values were lowest for early time points in both control and drought conditions, suggesting that predictions for these time point may be inaccurate. However, at later time points there was a high agreement between predicted and observed values for PSA. Collectively, these results suggest that the WSI-Gomp captures shoot growth trajectories in contrasting water regimes, however other factors not accounted for in the growth model also influence observed PSA values.

**Figure 3.**
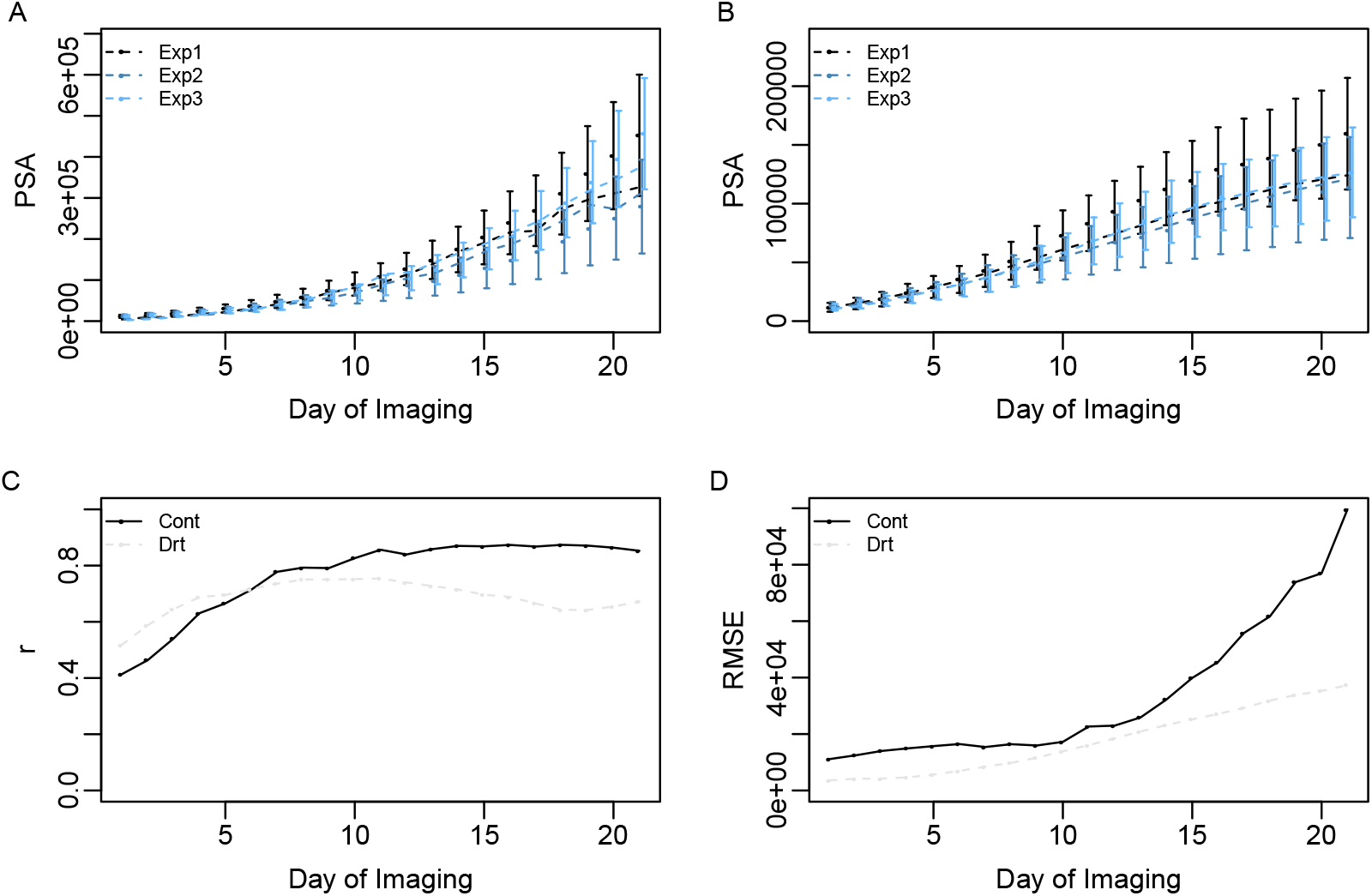
Capturing shoot growth trajectories using the WSI-Gomp model. Observed (points) and predicted (broken line) mean shoot growth trajectories for each experiment under control (A) and drought (B) conditions. For both A and B, Nelder-meader optimization was used to fit the WSI-Gomp model to the mean shoot growth trajectories for each experiment. (C) Average correlation between predicted trajectories and observed PSA values. (D) Root mean squared error between predicted trajectories and observed PSA values. For both C and D, the WSI-Gomp model was fit using the hierarchical Bayesian model and predicted PSA values were compared at each time point with observed values for each accession.

### Leveraging the growth model for biological inference

The advantage of the WSI-Gomp model is that it allows PSA trajectories to be modeled in response to declining soil water content, and provides a straight-forward way to calculate the predicted inflection point under an observed WSI value. With this in mind, we next sought to determine what observable characteristics influence the timing of this inflection point in drought conditions. To this end, we calculated the time of inflection (TOI) for each plant in drought by determining the earliest time in which the (*t* – *WSI^α^*) component of the model became positive (Supplemental File S2). As expected the predicted TOI were lower in drought conditions compared to control, indicating that the inflection of the growth curve occurs early under drought conditions compared to well-watered conditions (Figure 4A). TOI in drought-treated plants ranged from 8 - 16 days of imaging, while in control plants TOI values ranged from 14 - 20.

**Figure 4.**
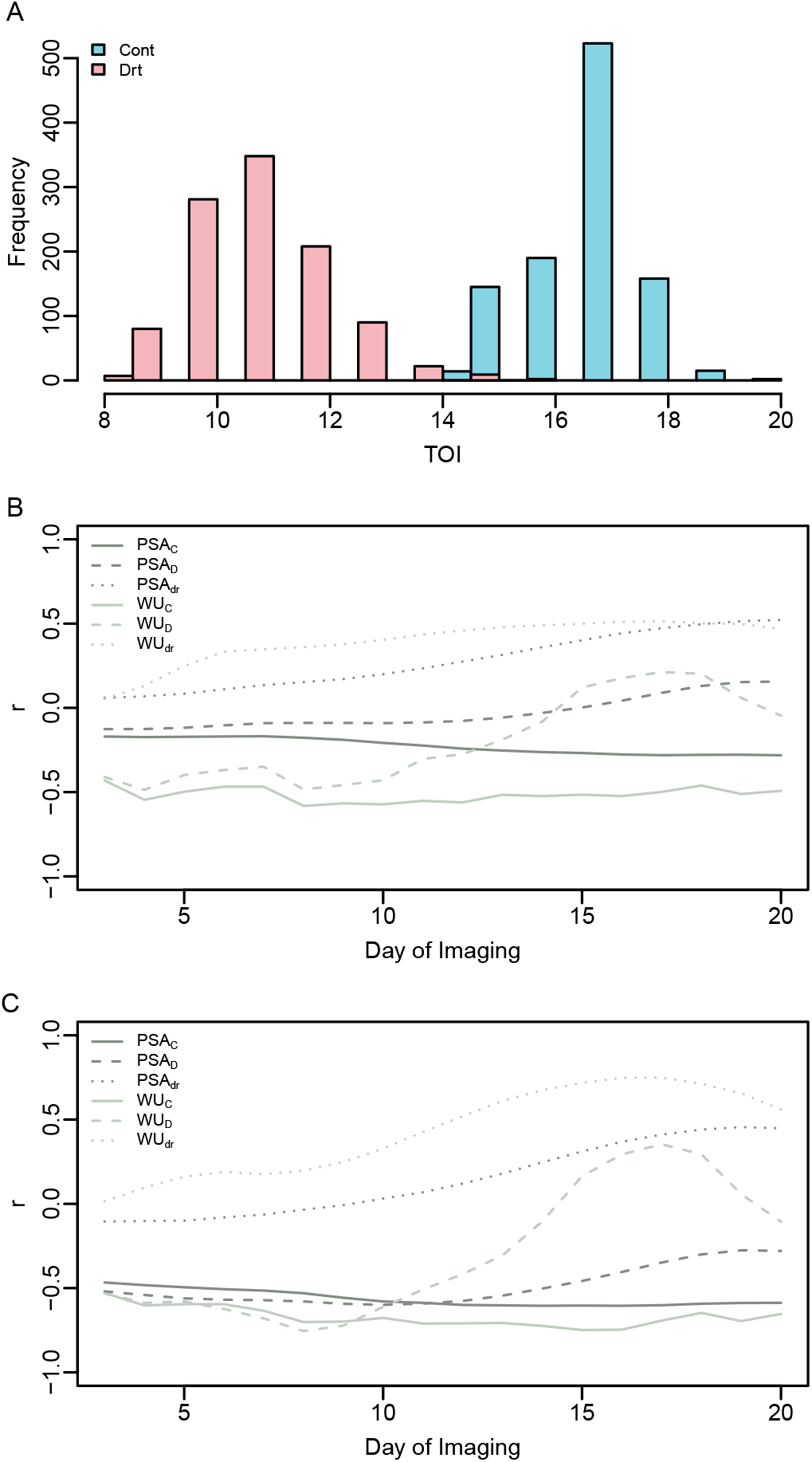
Distribution and interpretation of predicted time of infection. (A) The distribution of time of inflection (TOI) values in control and drought conditions. Correlation between time of inflection in control (B) and drought (C) and empirical observations for projected shoot area (PSA) and water use (WU). Spearman’s correlation was performed using a three-day sliding window.

To determine how observable phenotypes influenced TOI, the predicted TOI values were compared to water use (WU), PSA, and the ratio of these values in drought to control (indicated by the subscript “_dr_” meaning drought response) over the course of the experiment. Relationships were assessed using Spearman’s correlation using a sliding window of three-days (Figure 4B,C). In drought conditions, we observed a negative relationship between TOI in drought and PSA in both control and drought conditions, indicating that larger plants tend to have earlier retardation of shoot growth rate (Figure 4B). In drought, this relationship became weaker as the soil water declined and drought became more severe. This trend is likely because at these time points shoot growth in large plants were likely already repressed by drought. Similar, albeit slightly stronger, negative correlations were observed between WU in control conditions and TOI in drought (TOI_*D*_). An interesting trend was observed for WU in drought conditions and TOI_*D*_. At early time points (e.g. days 0-14) a negative correlation was observed between TOI_D_ and WU in drought. However, around day 15-18 this trend is reversed completely, with a positive correlation observed between WU and TOI_*D*_. As expected, TOI_*D*_ showed a positive relationship PSA_*dr*_ to drought (e.g. the ratio of PSA in drought to control), indicating that accessions with early inflection points tend to show a larger reduction in PSA under drought relative to control.

Similar trends were observed in control conditions, however the values of the correlation coefficients were different compared to drought (Figure 4C). A negative relationship was observed between TOI in control (TOI_*C*_) and PSA in control, which is consistent with the relationship observed for TOI_*D*_ conditions. However, TOI_*C*_ and PSA in drought showed a very weak relationship, showing a slight negative correlation during initial time points and a very weak positive relationship observed at later time points. Consistent with control conditions, the relationship between WU in control and TOI_*C*_ showed a strong positive correlation. Moreover, the correlation between TOI_*C*_ and WU in drought was negative at early time points and positive at later time points which is similar to the trend observed between TOI_*D*_ and WU. Although the interpretation of *α* and TOI is not very straightforward because plants were grown in the absence of water stress, the observed correlation suggests these parameters may have a similar interpretation as in drought conditions.

### Genome-wide association provides insight into loci influencing model parameters

Model parameter estimates for the WSI-Gomp model were obtained using a hierarchical Bayesian framework, wherein the growth model is fit in the first level and in the second level an EBL approach is used to predict marker effects from model parameters. Thus, the advantage of this approach is two-fold: first, solutions for model parameters are obtained by leveraging the genomic relationships among the accessions; secondly, the inferred marker effects can be used to identify genomic regions that influence the magnitude of the model parameters. Thus this information can be leveraged to identify QTL and potential candidate genes that may influence shoot growth trajectories in response to water deficit. To this end, we sought to utilize the inferred marker effects to identify genomic regions that regulate model parameters and influence dynamic shoot growth trajectories in response to water availability. The absolute value of inferred marker effects are provided in the Manhattan plots in Figure 5 and Supplemental File S3. Since obtaining *p*-values from Bayesian approaches is non-trivial, we report loci and candidate genes for the top-20 SNPs ranked based on the absolute value of marker effects (|β|).

**Figure 5.**
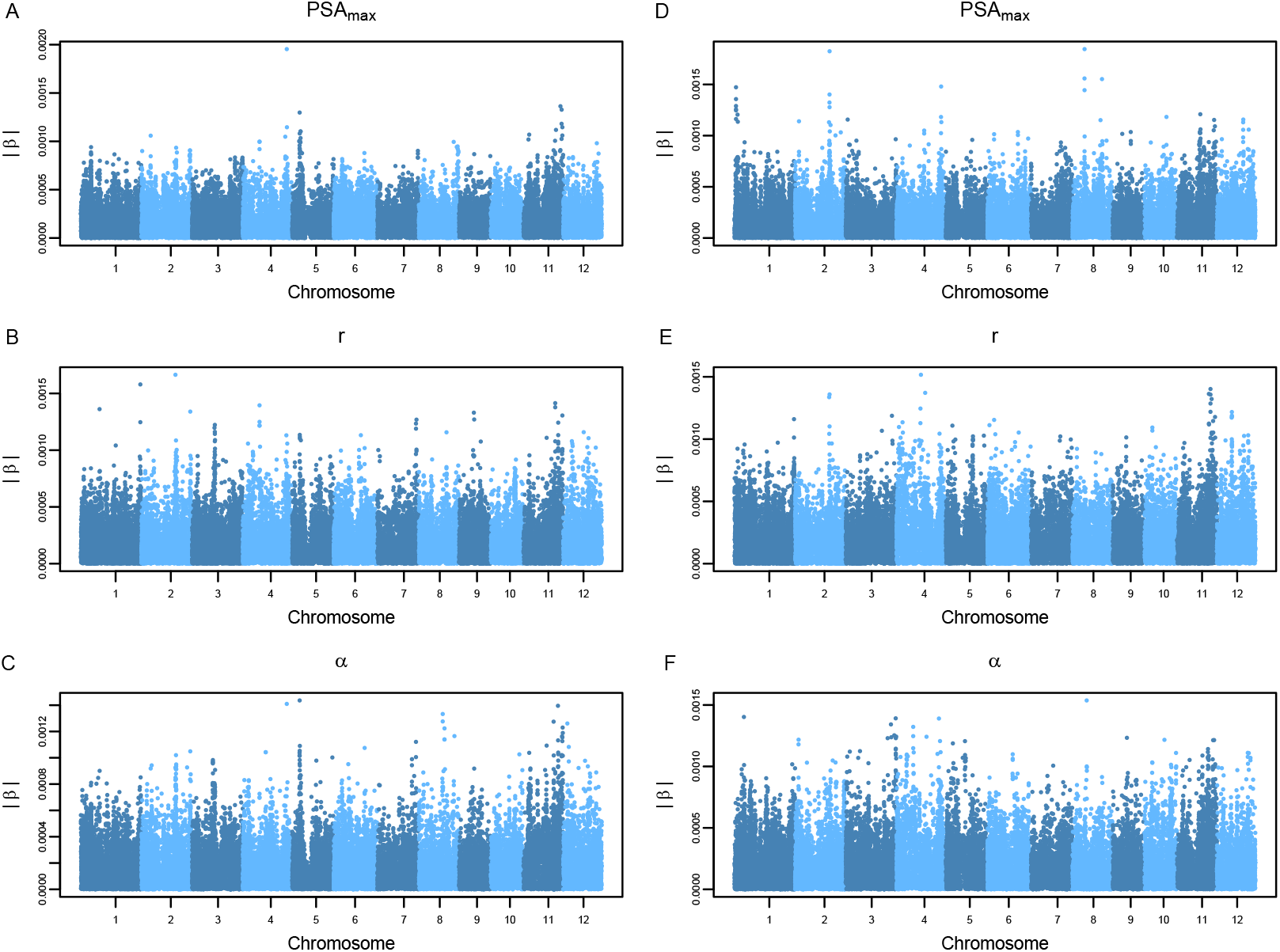
Genomic regions influencing model parameters. Predicted marker effects are shown for each of the WSI-Gomp model parameters. Panels A-C show marker effects for model parameters fit to growth trajectories in control conditions, while panels D-F show marker effects for drought conditions. The absolute value of predicted marker effects (|*β*|) is shown on the *y*-axis.

The model parameters *r* and *α* in both control and drought conditions exhibit a polygenic genetic architecture. We identified several markers with small contributions to the parameter values. Although the model parameters *α* and *r* showed a polygenic architecture, several notable genes were identified within the regions defined by SNPs with relatively larger effects (Supplemental File S4). For instance, at approximately 6.7 Mb on chromosome 1, a gene encoding an osmotin protein (*OSM34*) was found approximately 75 Kb upstream of the top SNP associated with *α* in drought within this region. Osmotin proteins play a role in plant biotic and abiotic stress responses, particularly drought stress (Narasimhan et al., 2009; Sharma et al., 2013). Additionally, a membrane-bound protein involved in chilling tolerance, *COLD1*, was found approximately 27 kB downstream of the SNP with the largest effect on chromosome 4 for *α* in drought (Ma et al., 2015). The presence of these two genes known to be involved in abiotic stress responses warrants further investigation.

The parameter *PSA_max_* showed a simpler genetic architecture in control and drought conditions. In control conditions, one large QTL was identified on chromosome 4 with the SNP with the largest effect located at approximately 31.4 Mb on chromosome 4. Within this region, a gene involved with the regulation of polar auxin transport, *Narrow Leaf1* (*NAL1*), was identified. Several studies have reported that variants in the *NAL* gene have pleiotropic effects and alter plant vascular patterning, spikelet number, leaf size, root system architecture, and shoot biomass (Qi et al., 2008; Fujita et al., 2013). In drought conditions, several QTL were identified for *PSA_max_*, with notable peaks located on chromosomes 1, 2, 4, and 8. The SNP with the largest effect was located at approximately 21 Mb on chromosome 8. Within this region, a gene known to regulate flowering time under short-day conditions was identified, *GF14c* (Purwestri et al., 2009). Moreover, a second gene known to influence biomass and seed size, *OsMPS* was identified on chromosome 2 at approximately 24.5 Mb (Schmidt et al., 2013). Since *PSA_max_* is a parameter that describes the maximum biomass for each accession, the presence of genes known to regulate flowering time and biomass is promising and suggests that this parameter is biologically meaningful.

### Elucidating the genetic loci influencing time of inflection in contrasting water regimes

As mentioned above, an advantage of the WSI-Gomp model is that it offers an intuitive framework to determine the time point at which an inflection in the growth curve occurs. Besides the parameters explicitly defined by the model, the time of inflection can also be considered an additional phenotype that can be analyzed using conventional genome-wide association mapping frameworks. With this in mind, we sought to identify QTL that were associated with the time of inflection using a Bayesian whole-genome regression approach (Supplemental File S3). Estimates for model parameters were combined with observed environmental covariates to solve for the TOI for each accession in drought and control conditions. Marker associations with TOI were assessed using a GWAS approach that accounted for the ordinal response variable, and results are discussed in the context of the top-20 ranked SNPs based on |*β*| (Figure 6).

**Figure 6.**
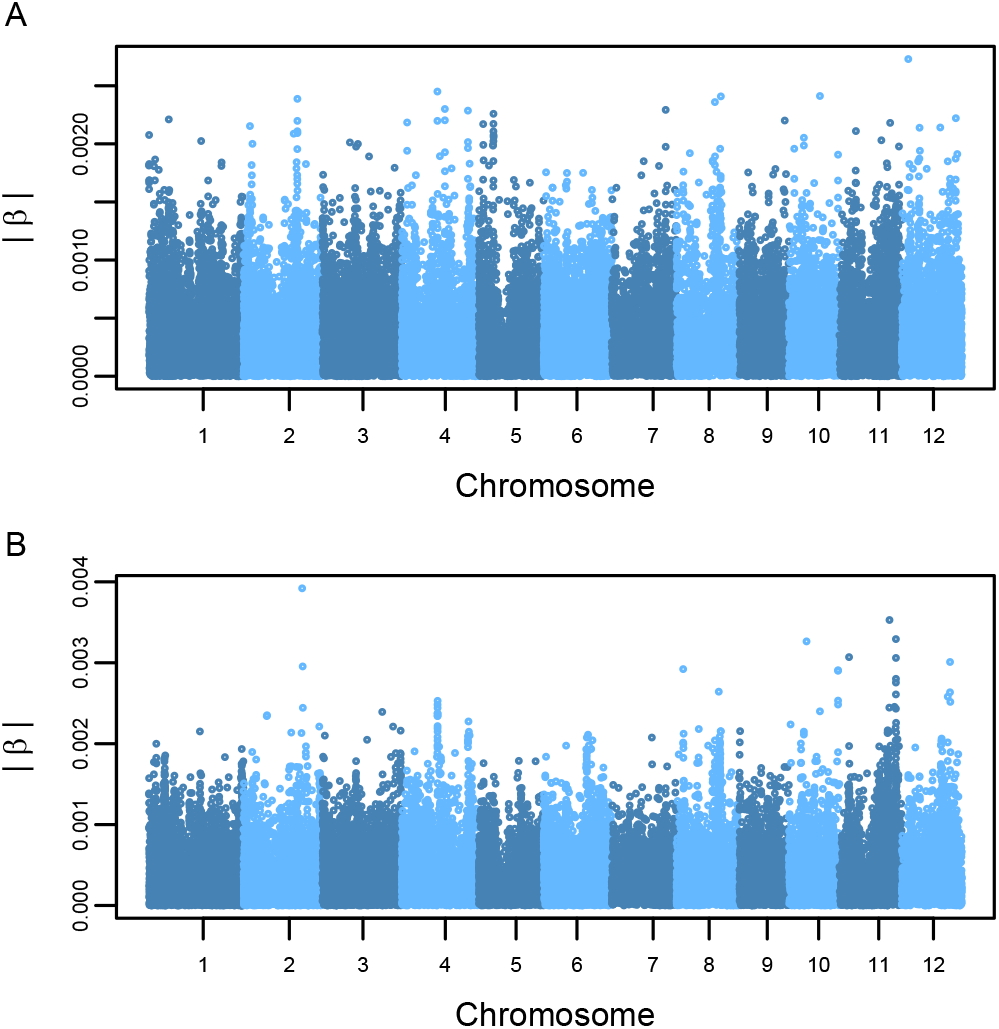
Manhattan plots for time of inflection (TOI) GWAS was conducted using TOI values in control (A) and drought conditions (B). Each point indicates a SNP marker and the *y*-axis shows the absolute value of predicted marker effects (|*β*|).

GWAS for TOI in control conditions showed that many SNPs have a small effect on the phenotype, indicating a complex genetic architecture for time of inflection in control conditions (Figure 6A). However for drought conditions, GWAS revealed two notable regions characterized by SNPs with relatively larger effects (Figure 6B). The first peak was identified at approximately 27 Mb on chromosome 2, while the second peak was located at 22.9 Mb on chromosome 11. Several notable genes were identified within these regions (Supplemental File S4). For instance, on chromosome 2 a gene encoding an aquaporin protein, *OsPIP1;1* was located approximately 674 bp from the second ranked SNP within this region (id2011870). Work by Liu et al. (2013) showed that *OsPIP1;1* functions as an active water channel and plays important role in salt tolerance, root hydraulic conductivity, and seed germination. The region on chromosome 11 harbored several genes known to be involved in disease resistance, however no genes had clear role in drought and/or abiotic stress tolerance. Thus, further studies are necessary to elucidate the role of this region in the drought responses.

To further explore the potential role of *PIP1;1* in influencing TOI under drought condition, we examined the expression of *PIP1;1* in shoot tissues of 87 diverse rice accessions grown in an ideal, controlled environment (Supplemental File S5). Gene expression levels were compared between major and minor allelic groups (*n* =74 and 13, respectively) using a linear model that accounted for population structure. This analysis showed significantly lower expression in accessions within the minor allelic group compared to the major allelic group (Figure 7). Moreover, accessions belonging to the minor allelic group also exhibited significantly higher TOI values, indicating that these accessions showed a later response to drought compared to those in the major allelic group. While additional studies are necessary to characterize the role of *PIP1;1* in influencing drought responses, large effect of this region on TOI in drought together with the differences in gene expression observed between allelic group are an encouraging direction for future studies.

**Figure 7.**
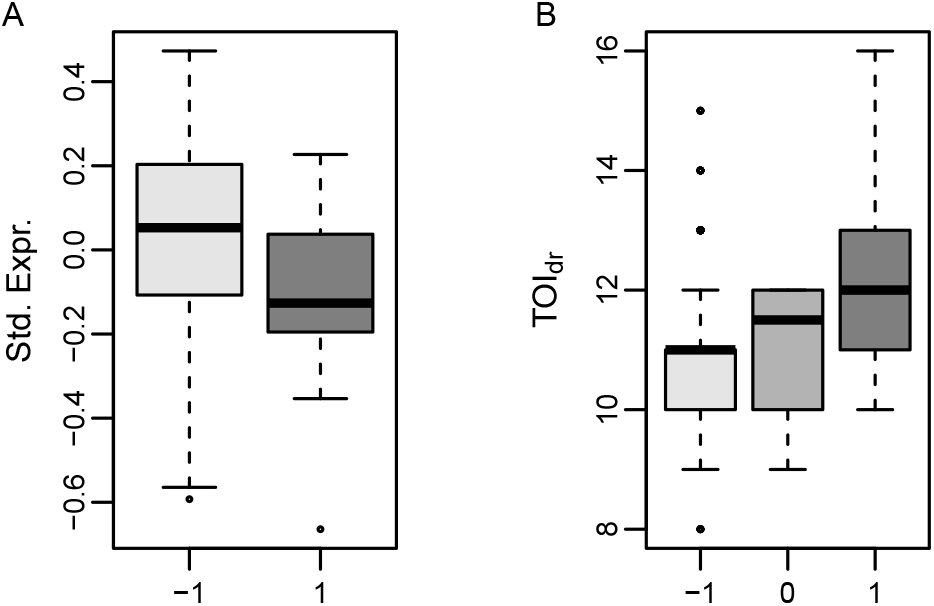
Comparing *OsPIP1;1* expression and TOI in drought between allelic groups at id2011870. (A) *OsPIP1;1* expression between minor (1) and major (−1) allelic groups at SNP id2011870. The values plotted in (A) are the residuals from a linear model that regressed standardized gene expression levels for *OsPIP1;1* on the top four principle components of the kinship matrix for 87 diverse rice accessions. Thus, these values represent the expression levels after correcting for population structure. (B) Comparison between allelic groups for time of inflection in drought conditions at SNP id2011870. The −1 corresponds to accessions that are homozygous for the major allele, 0 for those that are heterozygous, and 1 for accessions homozygous for the minor allele.

## Discussion

Drought tolerance during vegetative growth stage is most simply defined as the ability to maintain growth under water deficit. It is determined by the amount of water available to the plant and how efficiently is the water used to gain biomass. In terminal drought environments, where a fixed amount of water is available during the early season, the ability to maintain growth will be dependant on how well the plant can manage these resources through out the season. Thus, when studying drought tolerance, especially in terminal drought environments, it is important to jointly consider these factors. In the current study, we imposed a severe drought stress by completely withholding water for a period of 20 days (or until pots reached 20% FC). The effects of this severe stress was apparent soon after withholding water, as drought-stressed plants showed a significant reduction in shoot biomass after four days compared to control plants.

Given the importance of accounting for water availability when modeling temporal shoot growth trajectories, we developed a growth model that jointly models shoot biomass and soil water content. While the model parameters themselves can be used to describe characteristics of the growth curve and provide insight into the processes that influence shoot growth, the model can also be leveraged for additional biological inferences. For instance, we used genotype-specific parameter estimates to determine the point in which the growth rate begins to decline (i.e., TOI). Since the time values are standardized to be on the same scale as the WSI, this metric can be interpreted in two ways: (1) the time in which the growth rate begins to decline, or (2) the soil water content value that begins to repress growth. Regardless of the interpretation, this approach provides a means to assess drought sensitivity while accounting for variation in soil water content between plants.

### Joint modeling suggests a tradeoff between vigor and drought tolerance

Time of inflection provided biological insight into the relationship between plant size or vigor, and morphological responses to severe water deficit. Temporal correlation analyses between TOI and observed morphological and physiological responses revealed that large, vigorous plants tend to have an earlier decline in growth rate under severe drought conditions. Moreover, these plants tend to have high water demands in control conditions, and quickly exhaust soil water resources. The link between early vigor and drought responses has been studied extensively. Although some studies suggest that early vigor is advantageous in drought-prone environments, these benefits are highly dependant on the type of drought stress that is prevalent in these regions (Tardieu, 2011). A study by Kamoshita et al. (2004) evaluated six rice accessions under short and prolonged drought and examined the relationship between root system architecture, osmotic adjustment and biomass production. They found that highly vigorous accessions quickly developed a dense root system and extracted water quickly, but were also more sensitive to prolonged drought stress compared to low-vigor genotypes. However, these plants tended to recover more quickly after rewatering compared to low-vigor accessions. A more recent study by Rebolledo et al. (2012), found similar results and suggested that vigorous accessions also quickly exhaust starch reserves under prolonged drought resulting in a greater decline in biomass production compared to less vigorous accessions. Collectively, these studies support the observed negative correlation between plant size and drought sensitivity (as assessed with TOI), and suggests there is a trade-off between vigorous growth and the maintenance of growth in prolonged drought stress. Further studies are necessary to determine whether these relationships can be decoupled, or to identify the optimal balance between these two attributes.

### Leveraging the genome-enabled growth model for candidate gene discovery

The hierarchical Bayesian framework developed by Onogi et al. (2016) provides a powerful approach to improve the estimation of model parameters and to estimate the genomic contributions to the model parameters. Since the model parameters are regressed on genome-wide SNP markers, this framework can be used to calculate genetic values for model parameters thereby enabling genomic selection for certain growth curve characteristics, and thus provides a means to identify important loci that influence trait trajectories (i.e., GWAS). While the initial study by Onogi et al. (2016) showed both applications of the approach, their primary objective was genomic prediction. Here, we leveraged the genome-enabled growth modeling approach to identify genomic regions that influence dynamic drought responses.

Many of the model parameters show a complex genetic architecture characterized by many loci with small effects. However, several notable regions that exhibited relatively large effects were identified that harbored potential candidate genes. For instance, two notable peaks were identified on chromosomes 1 and 4 for the parameter *α* in drought conditions. Both regions harbored candidate genes that have been reported to regulate drought and/or osmotic stress responses in plants. The region on chromosome 4 harbored a gene that is known to regulate chilling tolerance in rice, *COLD1* (Ma et al., 2015). *COLD1* was shown to be involved with the Ca^+2^ signaling response to cold stress. In Arabidopsis, the *COLD1* orthologs, *GTG1* and *GTG2*, are membrane-bound ABA receptors (Pandey et al., 2006, 2009). However, *COLD1* exhibits GTPase activity that is absent in *GTG1/2* (Ma et al., 2015). Thus, further studies are necessary to determine whether *COLD1* participates in drought responses.

In addition to the candidate genes associated by model parameters, whole-genome regression performed with TOI in drought conditions revealed a potential role for additional genes in the genetic regulation of the timing of growth responses to drought. An aquaporin gene, *OsPIP1;1* was identified within a prominent peak on chromosome 2 associated with TOI in drought conditions. Aquaporins are a large family of proteins that were initially reported to act as water transporters, but have since been shown to also transport CO_2_ and H_2_O_2_ (Uehlein et al., 2003; Dynowski et al., 2008; Bienert and Chaumont, 2014; Maurel et al., 2015; Wang et al., 2016; Rodrigues et al., 2017). Aquaporins have received considerable attention as a potential target to modify whole plant water transport and improve water status during drought stress (Sadok and Sinclair, 2009; Devi et al., 2012; Choudhary and Sinclair, 2014; Schoppach et al., 2014; Grondin et al., 2016). Work by Grondin et al. (2016) showed that aquaporins account for approximately 85% of root hydraulic conductivity in rice under drought stress.

While the role of *OsPIP1;1* in drought tolerance remains to be elucidated, several studies have provided encouraging evidence that *OsPIP1;1* may play a role in mediating drought responses. First, work by Liu et al. (2013) showed that *OsPIP1;1* functions as a water channel and plays a role in seed germination and salt tolerance. Second, Grondin et al. (2016) showed that the expression of *PIP1;1* is induced by drought stress. Finally, work by Wu et al. (2014) showed that *OsPIP1;1* interacted with an leucine-rich repeat receptor-like kinase gene that regulates drought tolerance in rice. Moreover, Liu et al. (2013) showed that over-expression of *OsPIP1;1* increased root hydraulic conductivity, indicating that higher expression of *OsPIP1;1* increases water flux. In the current study, we examined gene expression levels for *OsPIP1;1* in 87 diverse rice accessions and found that accessions exhibiting lower expression of *OsPIP1;1* also exhibited later retardation in growth rate compared to accessions with higher expressions. Moreover, as stated above, we observed that plants with high water demands in control conditions tend to exhaust soil-water resources in water-limited conditions leading to an early retardation in growth rate (i.e. earier TOI). Although considerable work is necessary to establish a role of *OsPIP1;1* in the regulation of drought responses, the positive relationship between *OsPIP1;1* expression and root hydraulic conductivity as well as the observed relationship between *OsPIP1;1* expression and TOI provides an interesting foundation for future functional studies.

### Concluding remarks

Improving drought tolerance in rice is a challenging objective. Efforts to improve drought tolerance are hindered by the heterogenity in drought-prone environments, the breadth and complexity of traits underlying drought adaptation, and the difficulty in characterizing large populations for these traits. Recent advances in phenotyping technologies have provided an effective means to measure morpho-physiological traits frequently throughout the growing season, and provide plant breeders and geneticists with dense phenotypic data describing complex responses. However, these technological advances must be coupled with frameworks that accommodate these multidimensional data sets, while providing a means to leverage high density genotypic data to predict phenotypes and novel biological inference. In this context, the genome-enabled growth model proposed in a significant advancement towards addressing this need. The WSI-Gomp model provides a simple, biologically meaningful framework that can describe complex temporal responses using few parameters. Moreover, since genome-wide markers are used to estimate model parameters, the inferred marker effects can be used to study the genes that may contribute to these responses, estimate genetic values for model parameters for known individuals, as well as predict the phenotypes for new, uncharacterized individuals. This study is the first to leverage genome-enabled growth model for genomic inference in rice, and provides novel insights into the basis of dynamic growth responses to drought stress.

## Acknowledgments

Funding for this research was provided by the National Science Foundation (United States) through Award No. 1238125 to HW, and Award No. 1736192 to HW and GM.

## Supporting Information

- **Supplemental File S1:** Raw phenotypic data for all 349 accessions used to fit the Gomp-WSI model.
- **Supplemental File S2:** Model parameter and time of inflection estiamtes for all 349 accessions obtained from the Gomp-WSI model.
- **Supplemental File S3:** Marker effects for GWAS for model parameters and time of inflection.
- **Supplemental File S4:** Candidate genes for model parameters and time of inflection.
- **Supplemental File S5:** Best linear unbiased predictions (BLUPs) for gene expression in 87 accessions.

